# Genetic diversity varies with species traits and latitude in predatory soil arthropods (Myriapoda: Chilopoda)

**DOI:** 10.1101/2022.05.17.492264

**Authors:** D. K. Bharti, Pooja Yashwant Pawar, Gregory D. Edgecombe, Jahnavi Joshi

**Affiliations:** CSIR-Centre for Cellular and Molecular Biology, Uppal Road, Hyderabad, India; Natural History Museum, London, UK

**Keywords:** biogeography, centipedes, global study, life history, macrogenetics, mitochondrial COI

## Abstract

**Aim:** To investigate the drivers of intra-specific genetic diversity in centipedes, a group of ancient predatory soil arthropods.

**Location:** Global

**Time period:** Present

**Major taxa studied:** Centipedes (Class: Chilopoda)

**Methods:** We assembled a database of over 1200 mitochondrial cytochrome *c* oxidase subunit I sequences representing 120 centipede species from all five orders of Chilopoda. We used this sequence dataset to estimate genetic diversity for centipede species and compared its distribution with estimates from other arthropod groups. We studied the variation in centipede genetic diversity with species traits and biogeography using a beta regression framework, controlling for the effect of shared evolutionary history within a family.

**Results:** We observed a wide variation in genetic diversity across centipede species (0 to 0.1713), which falls towards the higher end of values among arthropods. Overall, 21.51% of the variation in mitochondrial COI genetic diversity in centipedes was explained by a combination of predictors related to life history and biogeography. Genetic diversity decreased with body size and latitudinal position of sampled localities, was greater in species showing maternal care and increased with geographic distance among conspecifics.

**Main conclusions:** Centipedes fall towards the higher end of genetic diversity among arthropods, which may be related to their long evolutionary history and low dispersal ability. In centipedes, the negative association of body size with genetic diversity may be mediated by its influence on local abundance or the influence of ecological strategy on long-term population history. Species with maternal care had higher genetic diversity, which goes against our expectations and needs further scrutiny. Hemispheric differences in genetic diversity can be due to historic climatic stability and lower seasonality in the southern hemisphere. Overall, we find that despite the differences in mean genetic diversity among animals, similar processes related to life history strategy and biogeography shape the variation within them.

## Introduction

Intra-specific genetic diversity (henceforth genetic diversity) is the amount of genetic variation present among individuals of a species and is an important component of biodiversity. It indicates the evolutionary potential of a species and is correlated with fitness and species’ response to environmental change (DeWoody et al., 2021). Genetic diversity can also have an influence on higher levels of biological organization by influencing species diversity, shaping communities (Vellend & Geber, 2005) and regulating ecosystem functioning (Raffard et al., 2019). Population genetic theory postulates that neutral genetic diversity increases with effective population size – the size of an idealized population that loses genetic diversity at the same rate as the observed population (Kimura, 1983), and mutation rate. A reduction in population size increases the sampling error in allele frequencies between generations, known as genetic drift, leading to the loss of genetic diversity (Charlesworth, 2009).

Previous studies have shown that genetic diversity is influenced by species traits and biogeography (Leigh et al., 2021). Species traits can modulate long-term effective population size by determining species’ responses to environmental fluctuations. On the other hand, biogeographic correlates determine the strength of environmental fluctuations experienced by species, and therefore can influence genetic diversity (Ellegren & Galtier, 2016). The strength of the relationship between species traits, biogeography and genetic diversity can be obscured by differences in mutation rates between lineages, which can vary based on the genetic locus under study (Nabholz et al., 2009).

Global-scale studies from well-studied taxa show that mitochondrial genetic diversity decreases with latitude, indicating a relationship between latitude and evolutionary rate or stability (Gratton et al., 2017, Manel et al., 2020, Miraldo et al., 2016). Global comparisons of nuclear genetic diversity reveal taxon-specific drivers of genetic diversity in animals, influenced by life-history strategy, environment, range size and position (De Kort et al., 2021). Taxon-specific studies show that traits indicative of life history strategy such as fecundity (Romiguier et al., 2014), reproductive mode (Paz et al., 2015) and body size (Mackintosh et al., 2019) are better predictors of genome-wide genetic diversity than census population size. Apart from life history, biogeographic variables related to range size and latitudinal position have been found to influence mitochondrial genetic diversity (Fujisawa et al., 2015). Both global-scale and taxon-specific studies have limited representation of arthropod groups, undersampling the richness of species traits, evolutionary history and ecosystems they offer. Additionally, arthropods vary widely in their genetic diversity, having some of the highest values of genetic diversity among animals (Leffler et al., 2012).

Among arthropods, the subphylum Myriapoda consisting of millipedes, pauropods, centipedes and symphylans (Fernández et al., 2018), has not been well-represented in global studies of genetic diversity, and macroecology studies in general (Beck & McCain, 2020; Thakur et al., 2020). The class Chilopoda has a 420 million year old evolutionary history and consists of over 3150 described species belonging to five orders (Edgecombe & Giribet, 2019). Centipedes are important venomous predators of the soil ecosystem and their taxonomic orders vary in their evolutionary age, diversity of families and species, and traits related to body size, vision, maternal care, habit (Edgecombe & Giribet, 2007) and venom composition (Jenner et al., 2019). Molecular markers, often in combination with morphological characters, have been widely employed in centipedes to uncover phylogenetic relationships, delimit species, identify cryptic species (Joshi & Karanth, 2012; Siriwut et al 2018; Wesener et al., 2015, 2016), and study the evolution of important species traits, such as blindness (Edgecombe et al., 2019; Vahtera et al., 2012) and maternal care (Fernández et al., 2014).

The variation in species traits among centipedes can potentially influence genetic diversity. Centipedes show a striking variation in body size (ranging from a few mm to up to 300 mm), which can influence genetic diversity by regulating local population abundance (White et al., 2007). Centipedes are predominantly sexually reproducing and show variation in their reproductive strategy, which can influence fecundity and long-term effective population size and thus genetic diversity (Ellegren & Galtier, 2016). While species from two orders (Scutigeromorpha and Lithobiomorpha) lay single eggs, others (Craterostigmomorpha, Scolopendromorpha and Geophilomorpha) brood multiple eggs and maternal care is also provided to hatchlings (Bonato & Minelli, 2002; Fernández et al., 2014). Another species trait that can influence genetic diversity through its association with habitat specialization or dispersal ability is blindness, seen in the order Geophilomorpha, in a few species of Lithobiomorpha, and in three families along with a few subterranean species within Scolopendromorpha (Edgecombe et al., 2019; Vahtera et al., 2012).

Given their low dispersal ability, the geographic distribution of centipedes is largely shaped by geological events and species vary widely in their latitudinal range and biogeographic affiliations (Bonato & Zapparoli, 2011; Edgecombe & Giribet, 2007; Joshi & Edgecombe, 2019; Joshi & Karanth, 2011; Joshi et al., 2020). In terms of range size, which may be correlated with abundance and thus genetic diversity, centipedes consist of island endemics such as *Craterostigmus* species (Vélez et al., 2012), narrow-range continental endemics such as *Ethmostigmus agasthyamalaiensis* (Joshi & Edgecombe, 2018) and *Rhysida sada* (Joshi et al., 2019), and species with cosmopolitan distributions such as *Pachymerium ferrugineum* or pantropical distributions like *Scolopendra morsitans* (Shelley et al. 2005). Their distribution patterns can be associated with traits related to reproduction (parthenogenesis in *Lamyctes emarginatus*; Andersson, 2006) or habitat specificity (human commensalism in *Scutigera coleoptrata*).

Despite their wide variation in species traits and biogeography, very few studies have documented genetic diversity and its geographic distribution in centipedes. A comparative study of two *Cryptops* species from the South Pacific indicated that the evolutionary age of species, rather than island size or isolation, determines genetic diversity and population genetic structure (Murienne et al., 2011). A study of an island endemic from Tasmania, *Craterostigmus tasmanianus*, showed the presence of significant population subdivision, which was correlated with geological divisions within the island (Vélez et al., 2012). Phylogeography of the circum-Mediterranean species *Scolopendra cingulata* suggests multiple colonization events from Aegean islands to the mainland since the Last Glacial Maximum, presence of relictual populations, and genetic differentiation in the mainland associated with geo-tectonic events dated to the Miocene (Oeyen et al., 2014; Simaiakis et al., 2012). Finally, *Digitipes coonoorensis* was shown to consist of monophyletic groups with strong population structuring across a biogeographic barrier in the Western Ghats, India (Joshi & Karanth, 2012).

There has been an increase in the representation of centipedes in publicly available sequence data in the last two decades, primarily arising from integrative taxonomic studies (Edgecombe & Giribet, 2019 and references therein) and regional barcoding efforts (eg. Spelda et al., 2011; Wesener et al., 2015). Among other genetic markers, the mitochondrial cytochrome *c* oxidase subunit I gene (COI), which is widely used as a DNA barcode, is well-represented across centipede species. The availability of global-scale publicly available sequence data for centipede species that vary with respect to their evolutionary age, species traits and biogeography motivated us to study their relationship with genetic diversity in a comparative framework. In this study, we specifically ask –

1. How is genetic diversity distributed across centipede species? We aimed to understand the range of genetic diversity seen in centipedes, an ancient soil arthropod clade with a 420 million-year evolutionary history, in the context of genetic diversity documented in other well-studied arthropod clades.
2. What are the species traits and biogeographic variables correlated with genetic diversity in centipedes? Based on theory, we expect to see a negative relationship between body size, maternal care (associated with low lifetime fecundity) and blindness (associated with habitat specificity and dispersal) relative to genetic diversity. These species traits can reduce effective population size leading to a reduction in genetic diversity. While latitudinal range is thought to be correlated with population size and may be positively associated with genetic diversity, the mean latitudinal position is expected to show the inverse relationship (Figure 1).

**Figure 1.**
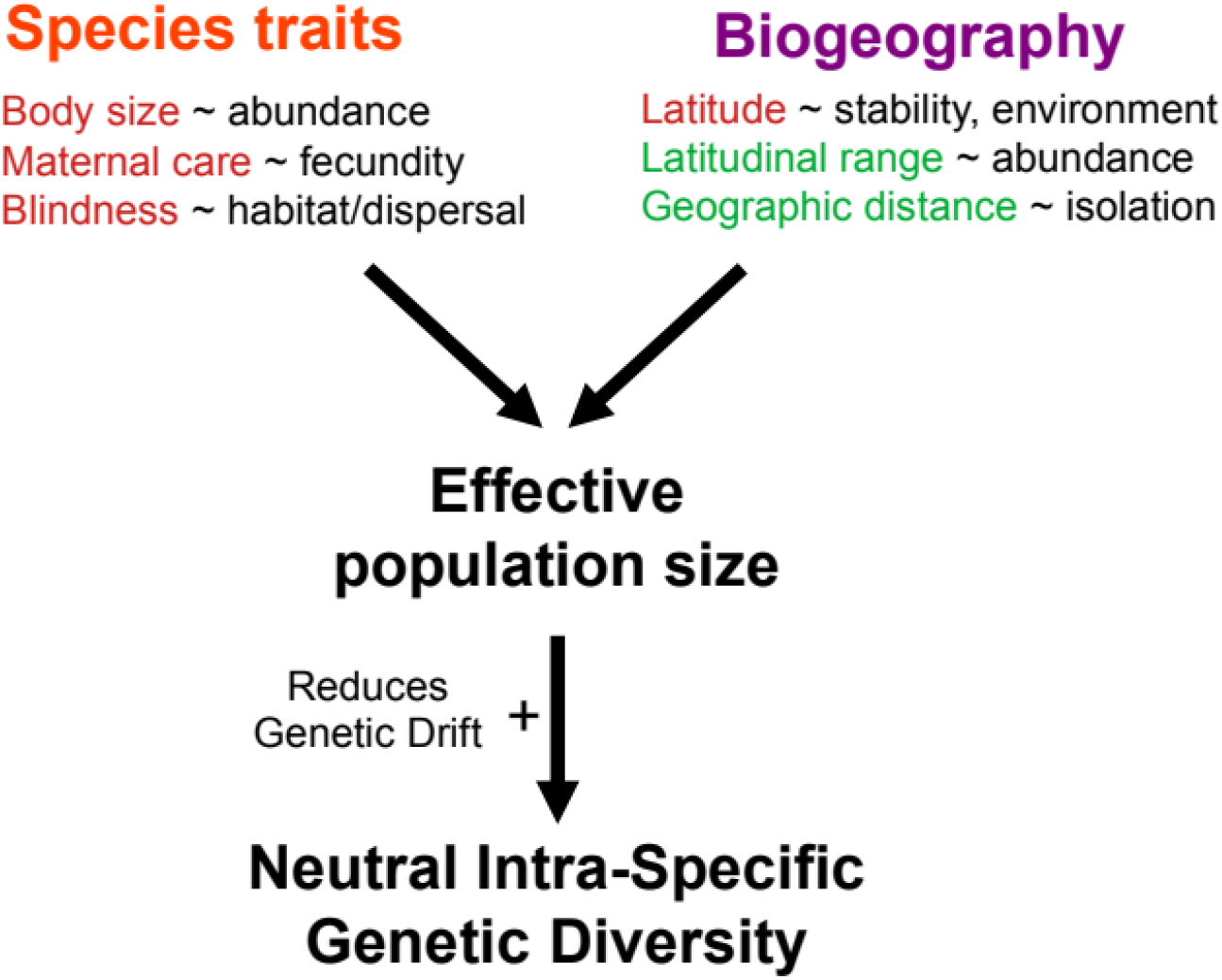
Schematic figure representing the theoretical drivers of intra-specific genetic diversity. Species traits and biogeography associated with species can influence their effective population size, which has a positive relationship with neutral genetic diversity. Variables with a negative influence on effective population size are highlighted in red and those with a positive influence in green.

A comprehensive global dataset including species traits, biogeographic correlates and mitochondrial sequences for 120 centipede species allowed us to estimate genetic diversity and examine its drivers. We observed a wide variation in genetic diversity across species, which was high compared to other arthropod classes. Both life history traits (body size and maternal care) and biogeographic correlates were important in explaining the variation in mitochondrial COI genetic diversity. This highlights the role of ecological strategy and latitudinal correlates of environmental stability as possible drivers of genetic diversity across living organisms, despite the differences in absolute values of genetic diversity between taxonomic groups.

## Material and methods

### DNA sequence data

We compiled sequence data from published studies and datasets and extracted accession numbers for the mitochondrial COI marker across the five centipede orders. In addition to accession numbers, we also compiled information on museum catalogue number, collection locality and geographic coordinates from source literature. We filtered this dataset to only retain those species that had at least three distinct sequence representatives (Appendix S1 in Supporting Information).

### Georeferencing sequences

Among the species that were retained, missing geographic coordinates associated with accession numbers were obtained by querying voucher numbers against museum websites (Appendix S1 in Supporting Information). When this was not available, we used geocoding to obtain geographic coordinates from locality names using the *geocode_OSM* function from the package ‘tmaptools’ (Tennekes, 2018). Average geographic distance between sequence locations for each species was calculated with the haversine formula using the function *geodist* function in the package ‘geodist’ (Padgham & Sumner, 2021) in R version 3.6.3 (R Core Team, 2021).

### Species traits and biogeographic information

Each species was supplemented with trait data from various sources. While the presence of maternal care and vision show variation at higher taxonomic levels (Edgecombe & Giribet, 2007), body size information for each species was obtained largely from species descriptions in taxonomic studies (Appendix S2 in Supporting Information). Species distribution information was collated from locations corresponding to species accession numbers, Chilobase 2.0 (Bonato et al., 2016), GBIF (GBIF.org, 2021), species descriptions and regional atlases. These distribution data were used to derive the latitudinal range for each species (Appendix S2 in Supporting Information). The mean latitudinal position of each species was calculated using only the geographic locations corresponding to the sequence dataset. We analysed two versions of the dataset to factor in native geographic ranges versus synanthropic introductions; one excludes six species that are likely introductions in parts of their sampled ranges while the other treats all records as potentially valid. The primary analysis described below was carried out using the smaller dataset representing the native range of centipede species.

### Sequence statistics

Mitochondrial COI sequences corresponding to the accession numbers were retrieved from the National Center for Biotechnology Information (NCBI) using the *entrez_fetch* function in the package ‘rentrez’ (Winter, 2017). For each species, sequence alignments were carried out separately using the MUSCLE algorithm in the package ‘muscle’ (Edgar, 2004) under the default parameters. The sequence alignment for each species was visualized in Aliview v1.26 (Larsson, 2014) and sequences were trimmed to bring them to the same length.

These edited alignments were used to calculate sequence statistics including sequence length, number of segregating sites (function *seg.sites* in the package ‘ape’; Paradis & Schliep, 2019), number of parsimony informative sites (function *pis* in the package ‘ips’; Heibl, 2008) and nucleotide diversity (function *nuc.div* in the package ‘pegas’; Paradis, 2010). Nucleotide diversity is calculated as the per site average number of differences between a pair of sequences, which is the sum of the number of differences between sequence pairs divided by the total number of sequence pairs compared. All analysis was carried out in R 3.6.1 (R Core Team, 2021).

### Statistical analysis

Genetic diversity is a proportion that estimates the probability of observing a mutation at a given site within a DNA sequence and can theoretically range from 0 to 1. However, intra-specific genetic diversity ranges closer to 0, as it is calculated from closely related individuals belonging to a single species. Our estimate of genetic diversity, average pairwise difference, is calculated by counting the number of mutations along a sequence that is hundreds of base pairs long. Given that genetic diversity is a proportion calculated using a large number of total counts (sequence length), it resembles continuous proportions, which can be analyzed using a beta regression framework (Douma & Weedon, 2019).

The error distribution of our regression model, the beta distribution, belongs to the exponential family and is defined by two parameters – mean and precision. In the deterministic part of our model, our response variable of genetic diversity is predicted by species traits (body size – continuous; blindness and maternal care – binary) and biogeography (latitudinal range, mean latitude and geographic distance – continuous). Since some of our genetic diversity estimates took zero values, which cannot be modelled using the beta regression algorithm, we replaced these with a small value following standard recommendations (Smithson & Verkuilen, 2006). All the predictor variables were standardized to have a mean of zero and standard deviation of 1. We used a logit-link function for the linear transformation of our exponential-family model. Our global model consisted of all the predictors mentioned above, where the precision parameter of the error distribution was independently modelled using sample size (the number of sequences representing each species), and taxonomic family was used as a random effect to account for the influence of shared evolutionary history on genetic diversity. Nested models created by dropping the precision parameter and/or the random effect were compared using their AIC values to choose the best model. We calculated a variation inflation factor for each predictor to check the influence of multi-collinearity between these variables on the coefficient estimates. We additionally measured the phylogenetic signal in model residuals using a family-level phylogenetic tree (Fernández et al., 2016) by calculating Pagel’s λ (Pagel, 1999). The scolopendromorph family Mimopidae represented by a single species *Mimops orientalis* was not represented in the phylogenetic tree and hence not included in calculating the phylogenetic signal.

The beta regression models were run using the *glmmTMB* function in the package ‘glmmTMB’ (Brooks et al., 2017) and the *betareg* function in the package ‘betareg’ (Cribari-Neto & Zeileis, 2010), and phylogenetic signal in model residuals was calculated using the *phylosig* function in the ‘phytools’ package (Revell, 2012) in R 3.6.1 (R Core Team, 2021)

## Results

### Geographic and taxonomic distribution of data

The complete dataset representing 46 published datasets with georeferenced sequences along with information on species trait and biogeography consisted of 1218 mitochondrial COI sequences representing 120 unique species, 12 of 18 centipede families and all five orders of Chilopoda. The species in our dataset varied in body size by two orders of magnitude (mean = 50 mm, range = 8.5 to 250 mm), with a relatively balanced distribution of reproductive strategy (78 of 120 species showing maternal care) and a predominance of species with the presence of vision (96 of 120 species). On average, each species was represented by 10 unique sequences (range = 3 to 68), with a mean alignment length of 653 bp (range = 465 to 840 bp). The centipede orders varied in the number of species and the total number of sequences representing them. These sequences arose from an average of eight unique geographic locations for each species (range = 1 to 53), separated by geographic distances up to 5066 km (Appendix S3 in Supporting Information).

Overall, the sequences in the dataset were obtained from 762 unique geographic locations spanning over 100 degrees in latitude (46.9°S to 60.5°N), with centipede orders showing distinct patterns of geographic distribution (Figure 2). For the two most well-represented orders, Scolopendromorpha sequences mostly originated from tropical and sub-tropical regions (mean latitude = 18.16°N), while Lithobiomorpha sequences were predominantly from northern temperate regions (mean latitude = 45.57°N). There were a larger number of sequences from the northern (n = 1042) as compared to the southern hemisphere (n = 176), with longitudinal under-representation from the Americas and Africa (Figure 2). These geographic gaps may arise from an interaction of differences in patterns of species distribution along with sequencing effort and taxon sampling.

**Figure 2.**
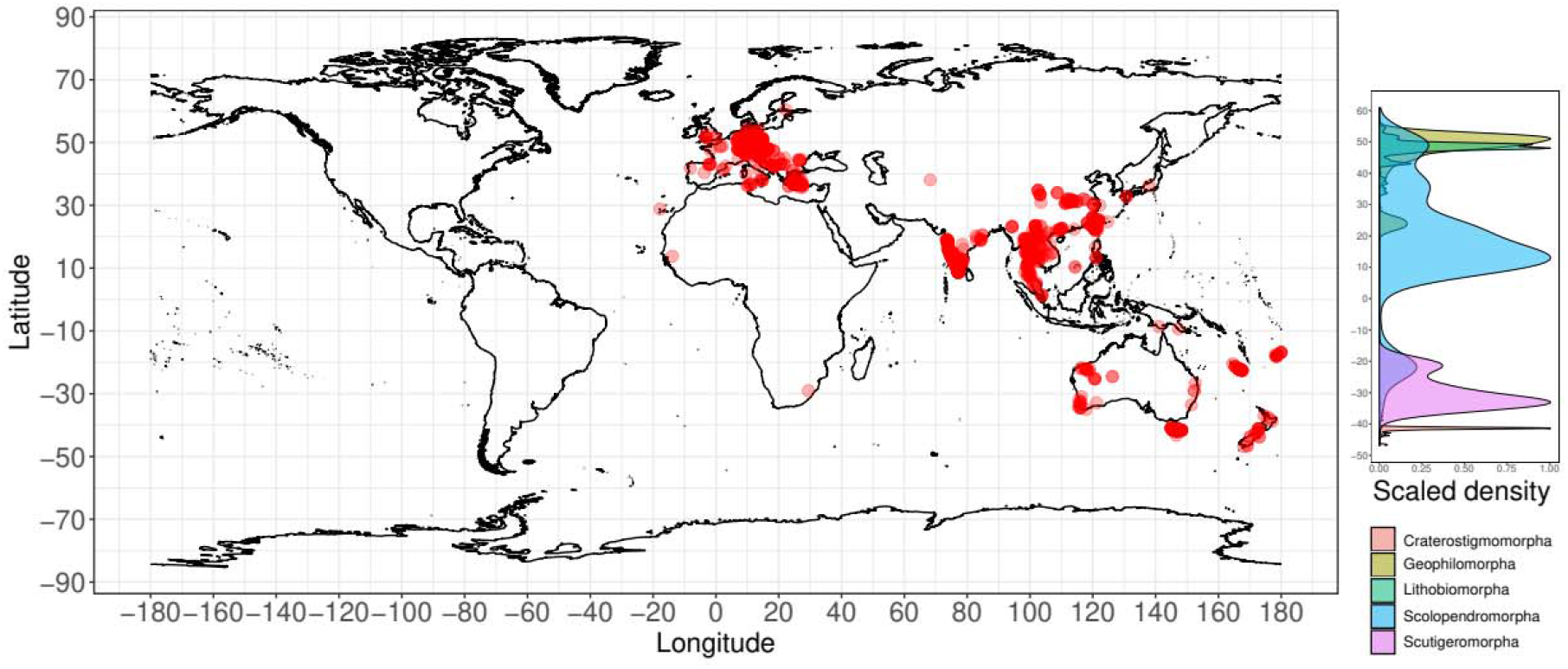
Global distribution of geographic locations associated with mitochondrial COI sequences used to calculate genetic diversity. The graph alongside shows the latitudinal distribution of locations based on taxonomic order, where the smoothed density of counts is scaled to a maximum of 1 for each order.

### Genetic diversity in centipedes compared to other arthropods

The average genetic diversity for centipedes was 0.0747 (range = 0 to 0.1713), with its distribution falling towards the higher end of values as compared to other arthropod groups (Figure 3). The average values of genetic diversity for other arthropod classes ranged from 0.0098 in insects to 0.0445 in millipedes, the latter belonging to the same sub-phylum as centipedes, Myriapoda. Though not an exhaustive effort, our collation of genetic diversity values from other arthropod groups showed evidence for an increased representation in insects in comparison to other taxonomic classes (Figure 3; Appendix S4 in Supporting Information).

**Figure 3.**
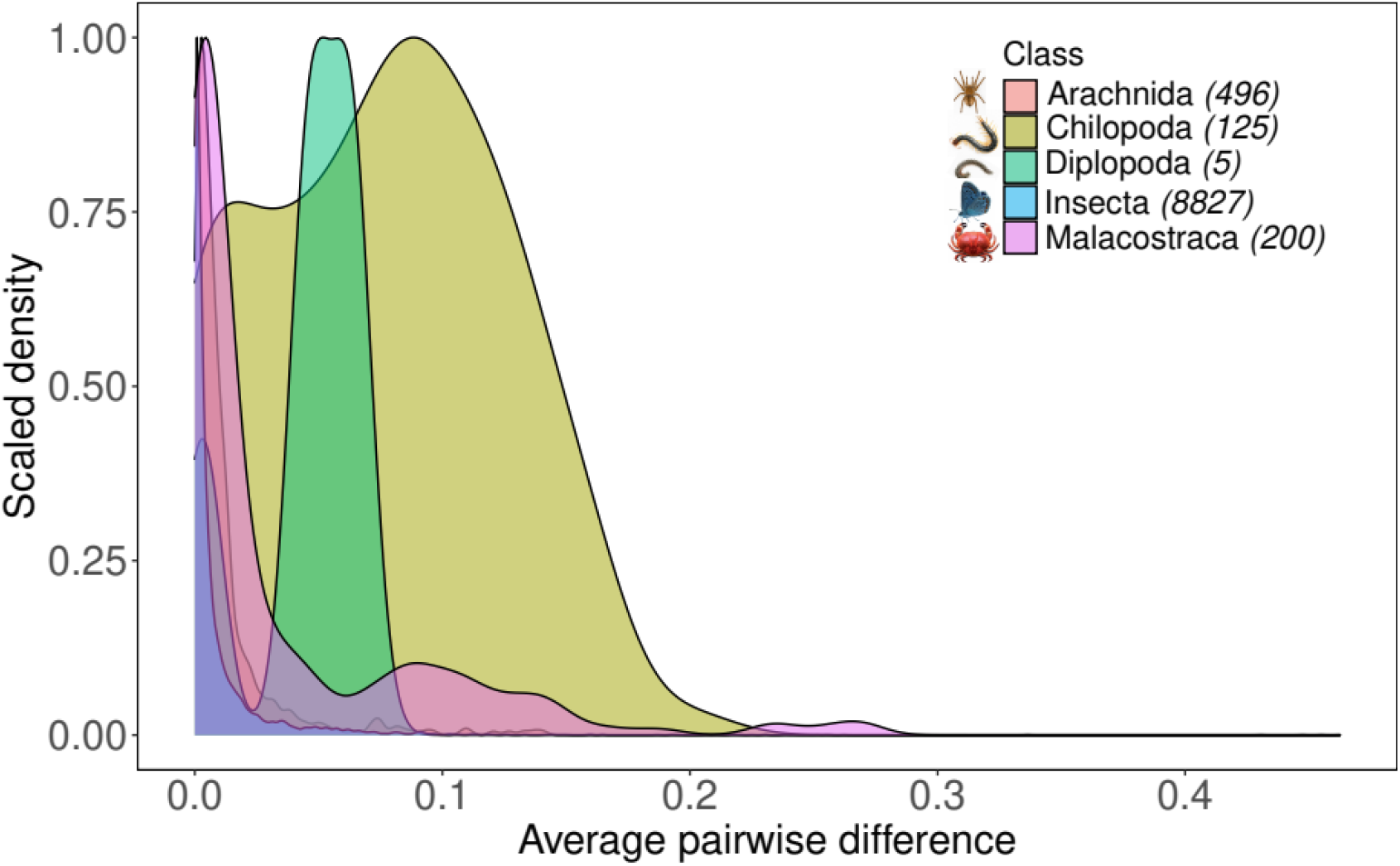
Distribution of mitochondrial COI genetic diversity (average pairwise differences) across arthropod groups. The values arise from the smoothed density of counts, scaled to a maximum of 1 for each taxonomic class. In the legend, the numbers in parentheses are the number of individual data points used for the taxonomic class. Data and their source references are provided in Appendix S4 in Supporting Information.

### Variation in genetic diversity is related to life history traits and mean latitude

Among the four models compared, the one using predictors for the fixed effects along with an independent predictor for precision emerged as the best model given its lowest AIC score (Table 1). This model explained 21.51% of the variation in genetic diversity across species and was selected over the global model, which used taxonomic family as a random effect. The variation inflation factors associated with the predictor variables were lower than 5, indicating that there was no significant influence of predictor multi-collinearity on coefficient estimates. Life history traits – body size and maternal care – significantly contributed to explaining this variation, while average geographic distance and the mean latitude of sequences were the biogeographic variables that emerged as significant (Table 1). The phylogenetic signal in the model residuals was close to zero and not significant (λ = 6.61 x 10^-5^, *p* = 1), indicating that the unexplained variation in genetic diversity could not be explained by a Brownian motion model of trait evolution at the family level.

**Table 1.** Parameter estimates from the best performing beta regression model defined as follows-

| Parameters | Estimate | 95% Confidence interval | z value |
| --- | --- | --- | --- |
| **Mean** |  |  |  |
| **Intercept** | -2.949***<br>(0.247) | -3.433– -2.466 | -11.959 |
| **Body size** | -0.230**<br>(0.078) | -0.383 – -0.077 | -2.949 |
| **Vision: Present** | 0.131<br>(0.176) | -0.214 – 0.476 | 0.743 |
| **Maternal care: Present** | 0.496**<br>(0.191) | 0.122 – 0.870 | 2.601 |
| **Mean latitude** | -0.211**<br>(0.066) | -0.340 – -0.082 | -3.213 |
| **Latitudinal range** | 0.072<br>(0.070) | -0.066 – 0.210 | 1.024 |
| **Average geographic distance** | 0.134*<br>(0.062) | 0.013 – 0.255 | 2.167 |
| **Precision** |  |  |  |
| **Intercept** | 3.271***<br>(0.133) | 3.010 – 3.531 | 24.633 |
| **Number of sequences** | 0.250*<br>(0.125) | 0.005 – 0.495 | 1.998 |
| **Pseudo R-squared** | 0.2151 |  |  |
| **Log-likelihood** | 213.7 |  |  |
| **AIC** | -409.4421 |  |  |
| **N** | 120 |  |  |
| **Significance** | *** = p < 0.001 |  |  |
|  | ** = p < 0.01 |  |  |
|  | * = p < 0.05 |  |  |

Genetic diversity showed a negative relationship with body size and mean latitude (Figure 4), where smaller species or those with sequences from lower latitudes had greater values of genetic diversity (Figure 5). Species with maternal care had higher values of genetic diversity, though it was associated with wide confidence intervals indicating substantial variation within each class (Figure 4, 5). Genetic diversity increased with greater average geographic distance between sequences, but the coefficient and its confidence interval were small in magnitude and close to zero. Confidence intervals of coefficients for vision and species latitudinal range overlapped with zero, indicating that they were relatively less important in explaining the variation in genetic diversity between centipede species (Figure 4).

**Figure 4.**
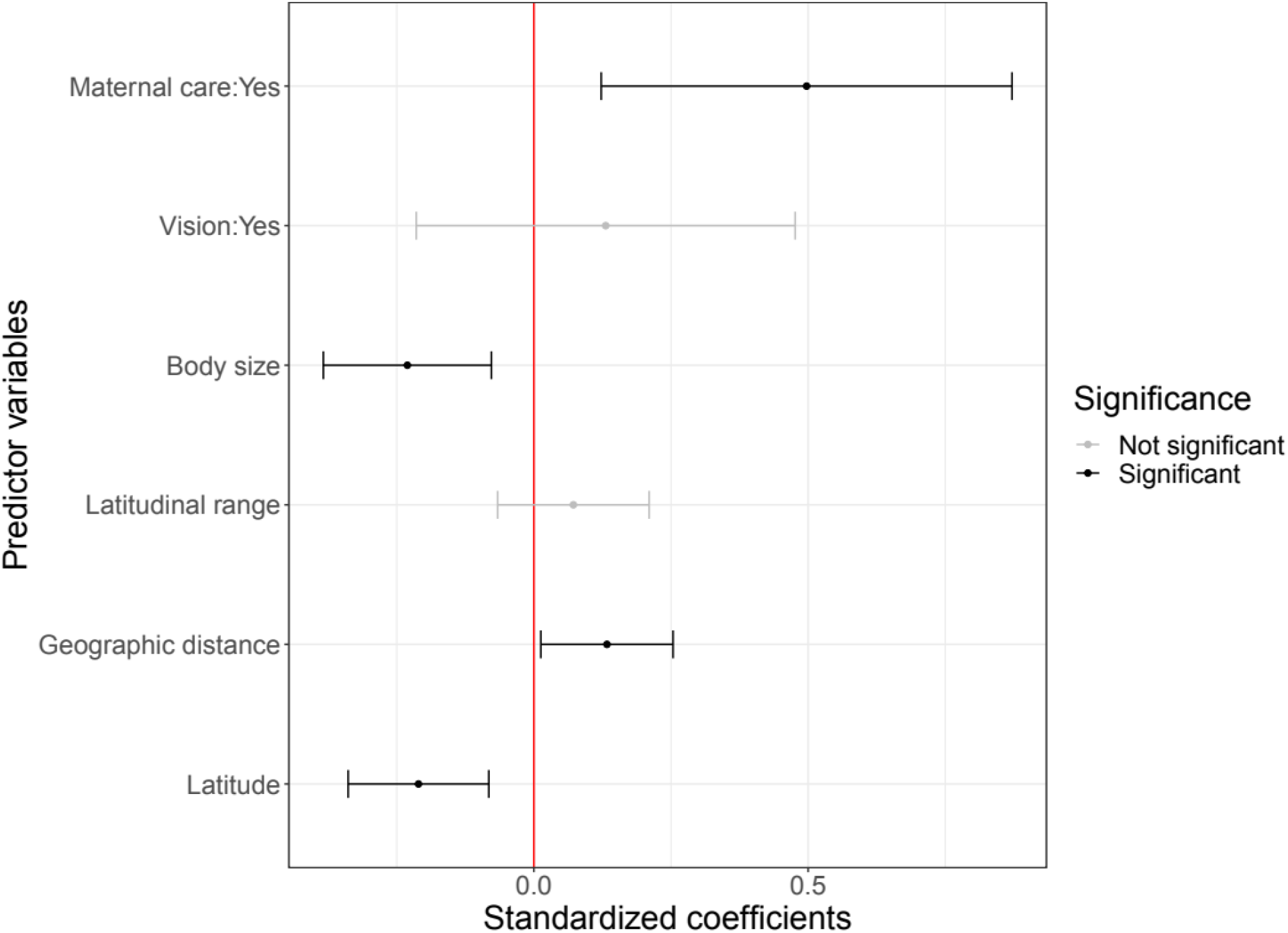
Standardized coefficient estimates (logit-scale) from the beta regression model with the lowest AIC value specified as - *Genetic Diversity ~ Body size + Vision + Maternal care + Mean latitude + Latitudinal range + Geographic distance / Number of sequences* Mean coefficient estimates are represented as points and their 95% confidence intervals are displayed as error bars for each predictor variable. Positive values indicate a positive relationship between the corresponding predictor variable and genetic diversity and the converse.

**Figure 5.**
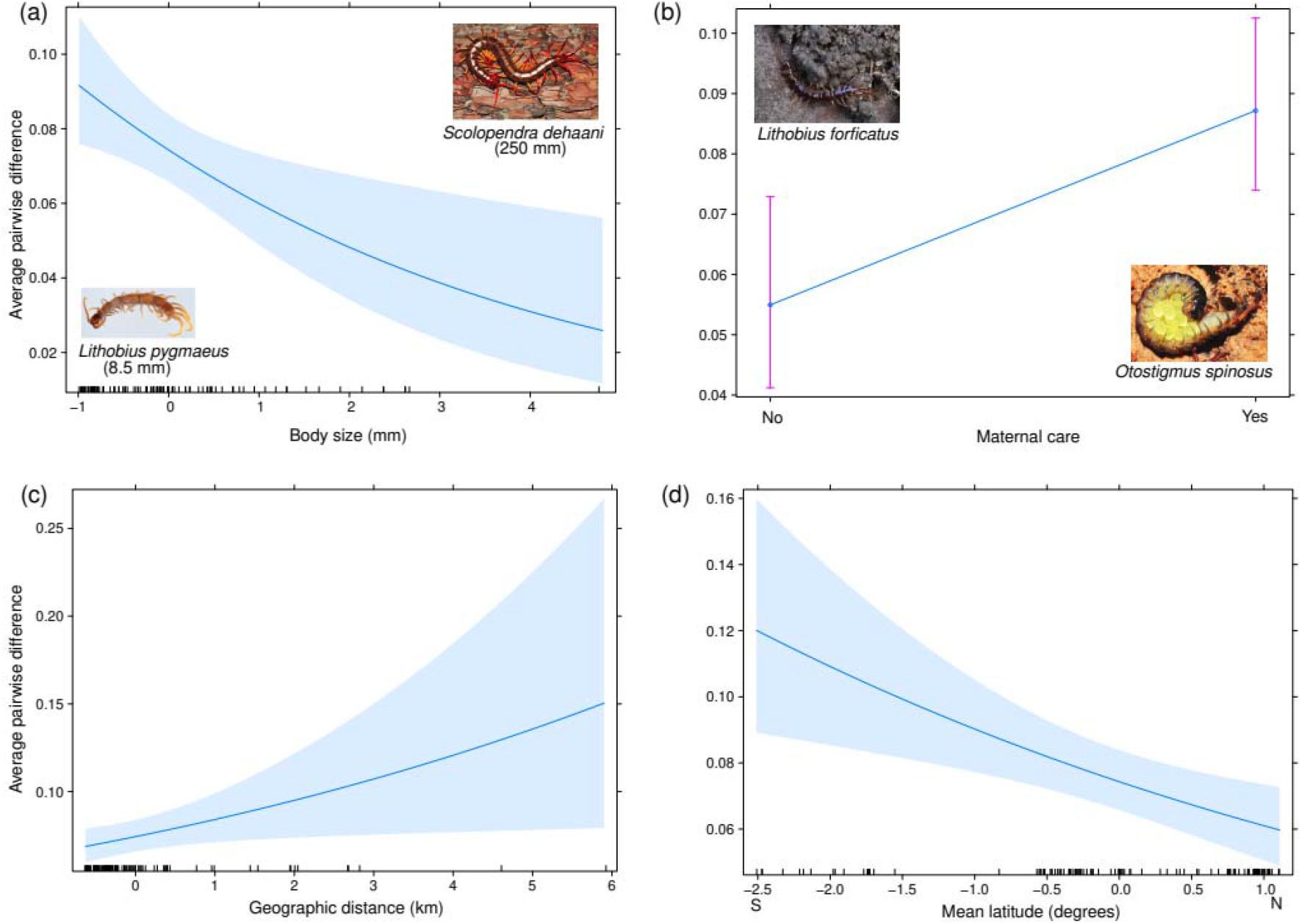
Fitted relationships between the significant explanatory variables and genetic diversity (measured as average pairwise difference) from the beta regression model in the scale of observed values. 95% confidence intervals are represented by the shaded band around the fitted line for continuous variables and error bars for the categorical variables. The effect of each predicted variable is calculated by varying it across the observed range, while keeping other predictor variables at their mean values.

In the analysis carried out using the dataset including likely synanthropic introductions, vision showed a significant positive relationship with genetic diversity, along with body size and maternal care. Values of average geographic distance showed a large variation between the two datasets and it was not a significant predictor when likely introductions were retained (Appendix S5 in Supporting Information).

## Discussion

### Centipedes have relatively high genetic diversity among arthropods

We find that centipedes have a high genetic diversity in comparison to other arthropod groups, which themselves fall in the higher end of the spectrum as compared to plants and chordates (Leffler et al., 2012). Among arthropods, where observations are skewed towards insects, high genetic diversity is hypothesized to be driven by their ability to reach large population sizes (Leffler et al., 2012). However, this mechanism may not hold true for centipedes, which are predatory arthropods that occur in low population densities in the soil ecosystem. The observed range of genetic diversity in centipedes may be explained by their persistence over a prolonged evolutionary history (‘evolutionary framework’ in Lawrence & Fraser, 2020) that extends back 420 million years (Edgecombe & Giribet, 2019). The limited dispersal ability of centipedes can also contribute to strong spatial differences in genetic composition reported in soil arthropod communities (Arribas et al., 2021) and the presence of geographically unique genetic diversity (Gloss et al, 2016). The positive relationship between geographic distance and genetic diversity seen in centipedes supports such a distance decay in genetic similarity. Additionally, it is possible that the presence of cryptic diversity may contribute to the relatively high values of genetic diversity observed for some species, which needs to be further examined with species delimitation methods using genome-level data.

### Species traits are significant correlates of genetic diversity

Despite the differences in the absolute values of genetic diversity across taxonomic groups, we find an overlap in the causal processes driving variation in genetic diversity. The genetic diversity of centipedes decreases with increasing body size, a relationship that has been observed across several animal groups (Brüniche-Olsen et al., 2021; De Kort et al., 2021; Mackintosh et al., 2019; Romiguier et al., 2014; an exception being Barrow et al., 2021). This association could be driven by the negative relationship between body size and abundance due to resource constraints (White et al., 2007), and by body size representing an ecological strategy that determines long-term effective population size (Ellegren & Galtier, 2016). Species with small body size, high fecundity and a short lifespan are hypothesized to recover from bottlenecks driven by environmental fluctuations more easily, therefore maintaining a larger long-term effective population size and greater genetic diversity (Ellegren & Galtier, 2016; Romiguier et al., 2014).

However, we find that centipede species showing maternal care of offspring had higher values of genetic diversity as compared to those that abandon their eggs. This questions our assumption of maternal care translating to greater investment in offspring quality over quantity, and therefore, lower lifetime fecundity and genetic diversity. There is a dearth of information on breeding biology from orders lacking maternal care, with substantial variation in the number of eggs reported for a few species (Lewis, 1981), and very little information on their survivorship. Gathering more natural history information would clarify the relationship between maternal care and lifetime fecundity in centipedes, and the observed positive relationship with genetic diversity.

The effect of blindness on genetic diversity may be mediated through its association with specialization to a subterranean habitat and/or low dispersal ability. We find that vision only emerges as a significant positive correlate of genetic diversity when synanthropic introductions are included, indicating sensitivity to changes in input data. While the observed pattern aligns with a negative association between specialization and genetic diversity seen in amphibians (De Kort et al., 2021), parasitoid wasps (Bunnefeld et al., 2018) and bumble bees (Jackson et al., 2018), other studies show no relationship in butterflies (Mackintosh et al., 2019), forest carabid beetles (Brouat et al., 2004) and bees (Dellicour et al., 2015). A more balanced representation of species with and without vision (e.g. more Geophilomorpha) using an expanded dataset could help resolve this relationship in centipedes.

### Latitudinal gradient and hemispheric differences in genetic diversity

Apart from species traits, several recent studies document a decline in genetic diversity with increasing latitude (beetles – Fujisawa et al., 2015; amphibians – Gratton et al., 2017; Miraldo et al., 2016; mammals – Millette et al., 2019; Theodoridis et al., 2020; salamanders – Barrow et al., 2021; amphibians and molluscs – De Kort et al., 2021), mirroring the latitudinal gradient in species diversity (Mittelbach et al., 2007). The mechanisms shaping latitudinal patterns in genetic diversity are thought to be congruent with those driving species diversity, related to climatic stability, longer evolutionary history, larger area with higher productivity and higher temperature resulting in high rates of molecular evolution at low latitudes (Fine et al., 2015). In centipedes, we find that genetic diversity increases from the northern hemisphere towards the tropics and the southern hemisphere. The hemispheric differences in genetic diversity of centipedes are indicative of similar patterns in species diversity (Dunn et al., 2009), which may be driven by differences in the current range of environmental variables and historic climatic stability (Chown et al., 2004). However, the wide confidence intervals for the southern hemisphere and limited representation of data points do not allow us to comment on a trend in comparison with the tropics.

Other arthropod groups have been reported to show deviations from the commonly observed latitudinal gradient in genetic diversity. Insects show a bimodal latitudinal distribution of genetic diversity, which may be related to the tropics having more specialized species with narrow niches and smaller ranges. On the other hand, the prevalence of diapause in the mid-latitudes may promote greater population persistence and genetic diversity in this region (French et al., 2022).

### What can intra-specific genetic diversity tell us about species diversity?

Variation in genetic diversity can be indicative of broader patterns in species diversity, either through the same underlying mechanisms acting independently or because of a cause-and-effect relationship between the two. As mentioned earlier, area, time, environmental factors and climatic stability can influence intra-specific and species diversity in parallel. Genetic diversity can positively influence species diversity if it reflects population fitness and reduces extinction rates or increases the diversity of competing species. High species diversity can negatively influence genetic diversity if species packing leads to niche specialization and if limiting resources result in smaller population sizes per species (Vellend & Geber, 2005).

In an empirical evaluation, neutral mechanisms involving area and isolation were found to be associated with both species and genetic diversity in beetles within an island system, shaping community and haplotype similarity along with dispersal ability (Papadopoulou et al., 2011). A strong relationship between genetic diversity and phylogenetic diversity was also observed in a global study of mammals, where it was speculated that microevolution at the population level may drive patterns in species diversity through various mechanisms (Theodoridis et al., 2020). However, the relationship between species and genetic diversity may be decoupled due to biological differences between taxa, lack of correlation between range size and genetic diversity (as opposed to a strong relationship between range size and species diversity) and sampling biases at the population-level (Lawrence & Fraser, 2020). It remains to be seen if these two hierarchical levels of biodiversity are correlated in centipedes, and if there is a causal link between genetic diversity as a species trait and diversification rates among various centipede groups.

### Significant variation in genetic diversity using a mitochondrial marker

As explained above, we find substantial variation in mitochondrial genetic diversity in centipedes, which is correlated with species traits, geographic distance and latitudinal distribution. This is in contrast with some previous studies, which find very limited variation in mitochondrial as compared to nuclear estimates (Bazin et al., 2006; Mackintosh et al., 2019) and no correlation with species life history traits (Dapporto et al., 2019). Estimates of genetic diversity can vary based on the properties of the genetic marker – mode of inheritance, ploidy (Berlin et al., 2007), length of the genetic map (Mackintosh et al., 2019) and mutation rate variation among taxa (Nabholz et al., 2009). The limited variation in mitochondrial genetic diversity and its lack of correlation with effective population size is ascribed to repeated selective sweeps and loss of diversity through genetic draft, given its maternal inheritance and smaller genome (Gillespie, 2001). For these reasons, the use of mitochondrial markers has been criticised despite the wide availability of sequences arising from barcoding efforts (Paz-Vinas et al., 2021).

In this context, it is interesting that we find significant variation in diversity estimates across centipede species, which is associated with species traits. The existence of this variation could be due to the smaller population sizes of predatory arthropods, which can dampen the frequency of selective sweeps as beneficial mutations are lost to genetic drift (Piganeonau & Eyre-Walker, 2009). The strength of selection also depends on the nature and spatial structure of genetic variation, which shapes genetic diversity (Leffler et al., 2012).

While the predictors in this study explain over a fifth of variation in genetic diversity, the strength of the observed correlation is congruent with other studies at a similar scale (Leigh et al., 2021). The reduced explanatory power could be due to spatial and temporal variation in drivers and population histories that cancel out at a broad spatial and taxonomic scale, the potential importance of environmental variables that are absent from the analysis, or the choice of the genetic marker as detailed above.

### Taxonomic and geographic gaps in sequencing efforts

Apart from revealing potential drivers of variation in genetic diversity, our dataset revealed taxonomic and distributional gaps in sequencing effort. There is a dearth of sufficient sequence information from the Americas and Africa, leading to a longitudinal bias in the available data (Figure 2). This also adds to a latitudinal gap in sampling, as most sequence data in the southern hemisphere are from Australia and New Zealand (Figure 2). There is also a sampling bias in the Palearctic, where most available sequences are from Europe (Figure 2).

The existing sequence information used in our analysis represents about 4% of existing species diversity and 12 of the 18 centipede families. Among the five centipede orders, the sampling gap in terms of species and family representation is the starkest for Geophilomorpha, where we have sampled 12 of over 1300 species (Appendix S3 in Supporting Information, Edgecombe & Giribet, 2007). This group is unique in terms of its habitat, being obligate soil-dwellers, as well as its feeding behaviour. Geophilomorphs feed using a greater degree of liquid suction than other centipedes, and use their mandibles to sweep or rasp food instead of chewing, which may potentially influence their prey resource base and population dynamics (Lewis, 1981). These geographic and taxonomic gaps can be the focus of future sampling efforts to reassess the current results and would also contribute to centipede phylogenetics and biogeography.

### Significance of examining intraspecific genetic diversity among divergent taxa

Our study generates hypotheses of drivers of genetic diversity in a relatively under-studied taxonomic group with a deep evolutionary history. These can be tested for their generality by using controlled comparisons of species with contrasting traits and distribution patterns and by screening additional nuclear markers. The generation of such hypotheses and efforts to test their validity provide means of understanding the generality of macroecological patterns across under-studied taxonomic groups (Beck & McCain, 2020) from unique and poorly explored habitats showing high biotic and abiotic variability (Thakur et al., 2020). While large-scale biogeographic studies in centipedes can be challenging due to a ‘species identification bottleneck’ reported in other arthropods (French et al., 2022), our study can act as a stepping-stone for future work. It also generates hypotheses for landscape-level studies exploring environmental and historical drivers of genetic diversity, and its relationship with population genetic structure (Salinas-Ivanenko & Múrria, 2021) and phylogenetic diversity (Bharti et al., 2021).

## Data Accessibility Statement

The raw data used for analysis are provided in Appendices S1-S3. R scripts for data analysis will be made available at the corresponding author’s github profile on publication.

## Supporting Information

**Appendix S1**: Mitochondrial COI accession numbers and associated location coordinates used for data analysis

**Appendix S2:** Centipede species traits and biogeographic variables used for data analysis

**Appendix S3:** Summary statistics of genetic diversity, sequence data and species traits across centipede species

**Appendix S4:** Genetic diversity estimates for arthropod classes

**Appendix S5:** Beta regression analysis results using a dataset that includes records corresponding to synanthropic introductions

## Acknowledgements

D. K. Bharti was supported during this study by a IndiaAlliance DBT Wellcome grant (IA/I/20/1/504919) to Jahnavi Joshi. Pooja Yashwant Pawar was supported by a start-up grant from CSIR-Centre for Cellular and Molecular Biology, Hyderabad to Jahnavi Joshi. We thank Dr. Nesrine Akkari, Prof. Zoltán Korsós and Prof. Pavel Stoev for help with centipede size and distribution data. We thank Dr. Rohit Naniwadekar for input regarding statistical analysis, and members of the Evol-Ecology lab at CSIR-Centre for Cellular and Molecular Biology, Hyderabad for suggestions that helped in improving the manuscript.

## Statement of authorship

DKB and JJ conceptualized this work with support from GE and PYP. All authors contributed equally to data curation. DKB carried out the methodology, formal analysis and visualization with support from JJ, GE and PYP. DKB wrote the first draft of the manuscript with support from JJ, GE and PYP, and all the authors reviewed and edited the manuscript. JJ acquired the funding for this study.

