## Appendix S3 for "Genetic diversity varies with species traits and latitude in predatory soil arthropods (Myriapoda: Chilopoda)"

**Table S3.1.** Summary of sequence statistics, taxonomic information, species traits and biogeographic variables in the compiled dataset presented as a mean value across species. For columns summarising numerical data, the mean is provided along with the range of observed values within parentheses.

|  | **Overall** | **Order-level classification** | | | | |
| --- | --- | --- | --- | --- | --- | --- |
|  |  | **Craterostigmomorpha** | **Geophilomorpha** | **Lithobiomorpha** | **Scolopendromorpha** | **Scutigeromorpha** |
| **No. families** | **12** | 1 | 3 | 2 | 5 | 1 |
| **No. species** | **120** | 2 | 12 | 38 | 64 | 4 |
| **Mean no. sequences per species** | **10.15**  **(3 – 68)** | 42.5  (17 - 68) | 6.25  (3 – 19) | 8.79  (3 – 26) | 11.09  (3 – 66) | 3.5  (3 – 5) |
| **Mean alignment length per species (bp)** | **652.87**  **(465 – 840)** | 750.5  (745 - 756) | 635.33  (535 – 658) | 651.95  (609 – 676) | 646.69  (465 – 840) | 764.25  (656 – 811) |
| **Mean no. unique locations per species** | **7.86**  **(1 – 53)** | 12  (10 – 14) | 5.83  (2 – 18) | 7.21  (1 – 20) | 8.81  (1 – 53) | 2.75  (2 – 4) |
| **Average pairwise difference per species** | **0.0747**  **(0 – 0.1713)** | 0.1131  (0.1064 – 0.1198) | 0.0725  (0 – 0.1483) | 0.0586  (0 – 0.1596) | 0.0838  (0 – 0.1713) | 0.0705  (0.0172 – 0.1335) |
| **Mean body size per species (mm)** | **50.22**  **(8.5 – 250)** | 43.5  (37 - 50) | 48.92  (20 – 95) | 17.33  (8.5 – 48) | 71.99  (20 – 250) | 21.75  (15 – 27) |
| **Maternal care: Present** | **78** | 2 | 12 | 0 | 64 | 0 |
| **Vision: Present** | **96** | 2 | 0 | 38 | 52 | 4 |
| **Mean latitudinal range per species (degrees)** | **16.32**  **(0 – 71.66)** | 7.38  (2.31 – 12.45) | 17.45  (3.40 – 27.45) | 18.17  (0 – 37.41) | 16.05  (0 – 71.66) | 4.17  (0.65 – 10.30) |
| **Mean latitude of sequence data per species**  **(degrees)** | **23.05**  **(-42.41 – 51.95)** | -41.93  (-42.41 – -41.44) | 43.76  (22.98 – 51.83) | 41.80  (-41.5 – 51.95) | 13.31  (-34.67 – 50.20) | -29.04  (-33.98 – -21.26) |
| **Mean geographic distance between sequences per species (km)** | **486.40**  **(0 – 5065.76)** | 250.54  (116.74 – 384.33) | 352.30  (133.67 – 786.71) | 294.08  (0 – 1988.20) | 655.57  (0 – 5065.76) | 126.95  (26.61 – 200.61) |
