## Appendix S5 for "Genetic diversity varies with species traits and latitude in predatory soil arthropods (Myriapoda: Chilopoda)"

**Appendix S5.** Details of the dataset including all centipede species records (native and synanthropic) and summary of beta-regression results

The complete dataset, including records representing likely synanthropic introductions, consisted of six additional species (total of 126 species) with changes to the number of sequences and/or species latitudinal range for seven species common to both the datasets. In this dataset, the largest change in predictor variables was seen in average geographic distance between sequences, ranging up to 9586 km, and species latitudinal range, ranging up to 103.22 degrees. In terms of longitudinal distribution of data, likely introductions of European species into North and South America were retained in this dataset, which were the only sequence representatives from these continents. With respect to species traits, there were 80 species showing maternal care and 99 species with vision. In total, this dataset represented sequences from 825 unique locations and the mean genetic diversity across species was comparable to the analysis with introduced species excluded, at a value of 0.0727 (range = 0 to 0.1713) (Appendix S5 in Supporting Information).

In the analysis carried out using this dataset, the model using fixed effects with a precision parameter had the lowest AIC score, explaining 16.04% variation in genetic diversity across species. However, while the nature of relationship of life history traits (body size and maternal care) and mean latitude with genetic diversity remained the same, vision emerged as a significant morphological trait, showing a positive relationship with genetic diversity. The average geographic distance between sequences was not a significant predictor.

**Table S4.2.** Parameter estimates from the best performing beta regression model -

Genetic diversity_i_ ~ Beta(*μ_i_, ϕ_i_*)

logit(*μ_i_*) = Body size*_i_* + Vision*_i_* + Maternal care*_i_* + Mean latitude*_i_* + Latitudinal range*_i_ +* Average geographic distance*_i_*

*ϕ_i_ ~* Number of sequences*_i_*

The input dataset retained sequences and range extent from likely synanthropic introductions for species.

| **Parameters** | **Estimate** | **Confidence interval** | **z value** |
| --- | --- | --- | --- |
| **Mean** | | | |
| **Intercept** | -3.332*** (0.244) | -3.810– -2.854 | -13.676 |
| **Body size** | -0.248** (0.080) | -0.404 – -0.092 | -3.122 |
| **Vision: Present** | 0.436*  (0.177) | 0.089 – 0.782 | 2.467 |
| **Maternal care: Present** | 0.699*** (0.189) | 0.328 – 1.069 | 3.699 |
| **Mean latitude** | -0.180** (0.066) | -0.309 – -0.051 | -2.730 |
| **Latitudinal range** | 0.014  (0.087) | -0.157 – 0.185 | 0.162 |
| **Average geographic distance** | 0.100  (0.083) | -0.063– 0.262 | 1.202 |
| **Precision** | | | |
| **Intercept** | -3.188*** (0.130) | -3.433 – -2.47 | 24.503 |
| **Number of sequences** | 0.262*  (0.121) | 0.025 – 0.499 | 2.163 |
| **Pseudo R-squared** | 0.1604 | | |
| **Log-likelihood** | 222.431 | | |
| **AIC** | -426.8619 | | |
| **N** | 126 | | |
| **Significance** | *** = p < 0.001  ** = p < 0.01  * = p < 0.05 | | |

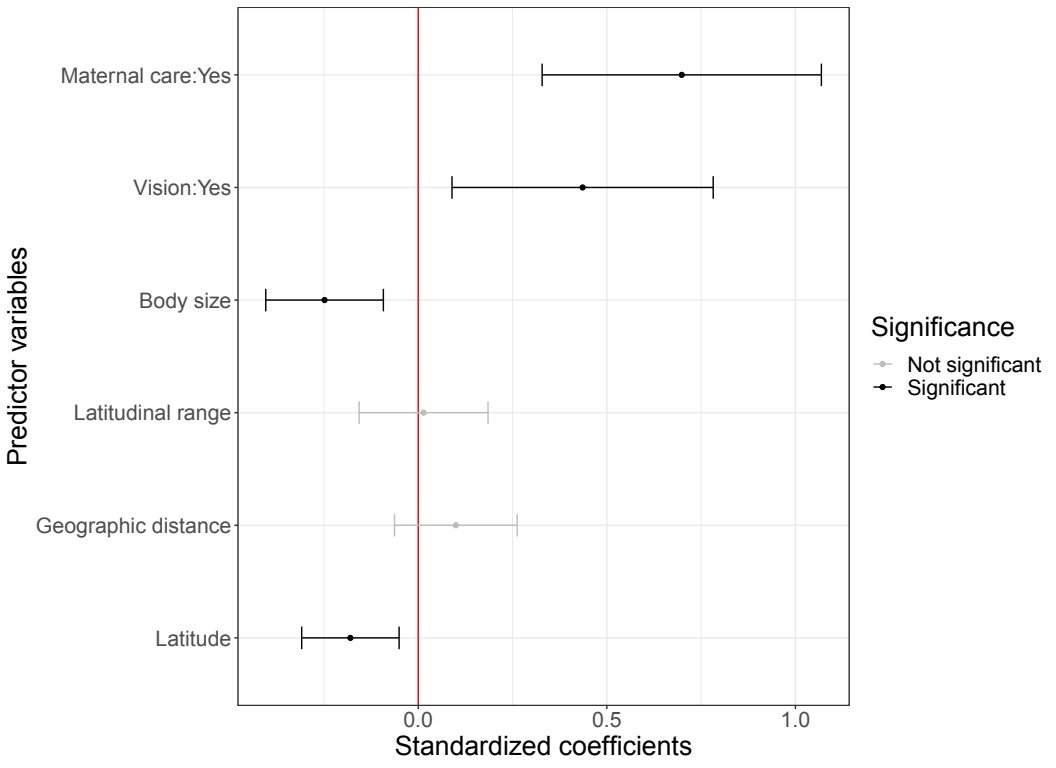

**Figure S4.1.** Standardized coefficient estimates (logit-scale) from the beta regression model with the lowest AIC value specified as -

*Genetic Diversity ~ Body size + Vision + Maternal care + Mean latitude + Latitudinal range* + *Geographic distance | Number of sequences*

The input dataset retains sequences and range extent corresponding to potential synanthropic introductions. Mean coefficient estimates are represented as points and their 95% confidence intervals are displayed as error bars for each predictor variable. Positive values indicate a positive relationship between the corresponding predictor variable and genetic diversity and the converse.
